# Modeling Glioma Intratumoral Heterogeneity with Primary Human Neural Stem and Progenitor Cells

**DOI:** 10.1101/2024.10.20.619254

**Authors:** Daniel Gao, Daniel Dan Liu, Anna E. Eastman, Nicole L. Womack, Benjamin F. Ohene-Gambill, Michelle Baez, Irving L. Weissman

## Abstract

Glioblastoma multiforme (GBM) is a deadly form of glioma notable for its significant intratumoral heterogeneity, which is believed to drive therapy resistance. GBM has been observed to mimic a neural stem cell hierarchy reminiscent of normal brain development. However, it is still unclear how cell-of-origin shapes intratumoral heterogeneity. Here, we develop a model of glioma initiation using neural stem and progenitor cells (NSPCs) purified from fetal human brain tissue. We previously described a method to prospectively isolate and culture tripotent neural stem cells (NSCs), bipotent glial progenitor cells (GPCs), and unipotent oligodendrocyte precursor cells (OPCs). We transduced these isogenic lines with dominant-negative TP53^R175H^ and NF1 knockdown, a commonly-used genetic model of GBM in mice. These reprogrammed lines robustly engrafted when transplanted into the brains of immunodeficient mice, and showed significant expansion over time. Engrafted cells were reextracted from the mouse brain for single cell RNA sequencing (scRNA-seq), in order to quantify how the cell-of-origin modulates the cellular subtypes found in the resulting tumor. This result revealed the strong influence the cell-of-origin plays in glioma heterogeneity. Our platform is highly adaptable and allows for modular and systematic interrogation of how cell-of-origin shape the tumor landscape.

## INTRODUCTION

Glioblastoma multiforme (GBM) is a deadly brain cancer with a median survival rate of 15 months^1^. Despite advancements in chemotherapy, immunotherapy and surgical techniques, GBM prognosis has remained poor. From 2001 to 2007, the one year survival rate improved meagerly from 29% to 39%^2^, and the three year survival rate improved only from 4.4% to 10%^3^. Much of this poor prognosis likely arises from the cellular heterogeneity within GBM tumors, which is believed to confer therapy resistance^4,5^.

Previous single cell transcriptomic studies on primary GBM tumors have identified distinct cellular subtypes that coexist within tumors^6^. Neftel et al. categorized four such transcriptomic “meta-modules”: the first consisted of genes involved in hypoxia, stress, or glycolytic function, and was dubbed mesenchymal-like. The other three meta-modules consisted of genes involved in neurodevelopment and were distinguished by their transcriptomic resemblance to neurons, astrocytes, or oligodendrocyte precursor cells (OPCs), and thus were called “neuron progenitor-like”, “astrocyte progenitor-like”, and “OPC-like” respectively. GBM tumors consist of an admixture of these four cellular subtypes at varying proportions. The presence of certain mutations has also been found to favor specific GBM subtypes, with EGFR mutations correlated with the astrocyte-like program, CDK4 mutations with the neuron progenitor-like program, PDGFRA mutations with the OPC-like program, and NF1 mutations with the mesenchymal-like program^6,7^. Other single cell transcriptomic studies on GBM have also proposed alternative classification schemes based on immune and metabolic gene signatures, which may be valuable from a more therapeutic perspective^8,9^. Interestingly, intratumoral heterogeneity appears to be biased based on the type of glioma. For example, while IDH-mutant oligodendrogliomas are enriched for differentiated oligodendrocyte-like tumor cells, H3K27M gliomas are predominated by proliferative OPC-like cells^10,11^.

During the development of the human cerebral cortex, radial glia residing in the ventricular zone serve as multipotent neural stem cells (NSCs), giving rise to the three major neural lineages: neurons, astrocytes, and oligodendrocytes. In humans, a second pool of radial glia known as outer radial glia (oRG) occupy a layer basal to the ventricular radial glia (vRG) and serve as an additional pool of neural stem cells. The admixture of various neural-like lineages within GBM tumors suggests the presence of a common progenitor, sometimes referred to as a GBM stem cell (GSC), though functional definitions vary^12–14^. Indeed, reference mapping of GBM cells onto normal fetal human brain cells suggests that GBM conserves a tri-lineage stem cell hierarchy^15^. GBMs also seem to harbor a population of cells resembling outer radial glia (oRG), which undergo characteristic mitotic somatic translocations and are suggested to contribute to GBM heterogeneity^16^. That glioma tumors revert into a fetal developmental ecosystem suggests that fetal NSPCs may serve as a useful system for modeling glioma heterogeneity and progression. In a similar vein, previous work has reprogrammed normal human lung and prostate epithelial cells into small cell lung and prostate cancer lines, and found that these reprogrammed cancer lines share a similar transcriptomic profile as patient-derived small cell lung and prostate tumors^17^.

Recently, we described a method to purify ten distinct neural stem and progenitor cell (NSPC) types from the fetal human brain^18^. These included the tri-potent vRG and oRG that give rise to neurons, astrocytes, and oligodendrocytes, a bipotent glial progenitor cell (GPC) that gives rise only to astrocytes and oligodendrocytes, and a unipotent oligodendrocyte progenitor cells (OPC) that gives rise exclusively to oligodendrocytes^18,19^. Using this prospective isolation strategy, we sought to investigate the role of cell-of-origin on glioma heterogeneity. Here, we develop an adaptable platform for reprogramming purified NSPC populations into glioma-like lines, demonstrating that these transformed lines recapitulate intratumoral heterogeneity *in vivo*.

## RESULTS

### Generation of isogenic NSPC lines for glioma modeling

Purified ventricular radial glia (vRG, henceforth NSCs), glial progenitor cells (GPCs), and pre-oligodendrocyte precursor cells (pOPCs) were isolated from fetal human brain tissue using FACS using our previously described purification strategy **(Figure 1A, B)**^20^. Briefly, following exclusion of debris, doublets, and non-neural lineages (Lin), NSCs were sorted as Lin^−^CD24^−^THY1^−/lo^EGFR^+^CXCR4^−^, GPCs as Lin^−^THY1^hi^EGFR^hi^PDGFRA^−^, and pOPCs as Lin^−^THY1^hi^EGFR^+^PDGFRA^+^. Each isogenic line was expanded in fetal growth media containing EGF, FGF, and LIF, and then transduced with lentiviral constructs carrying a dominant-negative TP53^R175H^ tagged with mCherry and shRNA against NF1 (shNF1) marked with mKOk, an established model of glioma induction **(Figure 1C)**^21^. NSPCs that received both lentiviral constructs were purified via FACS for mCherry and mKOk double positivity **(Figure 1D)**.

**Figure 1:**
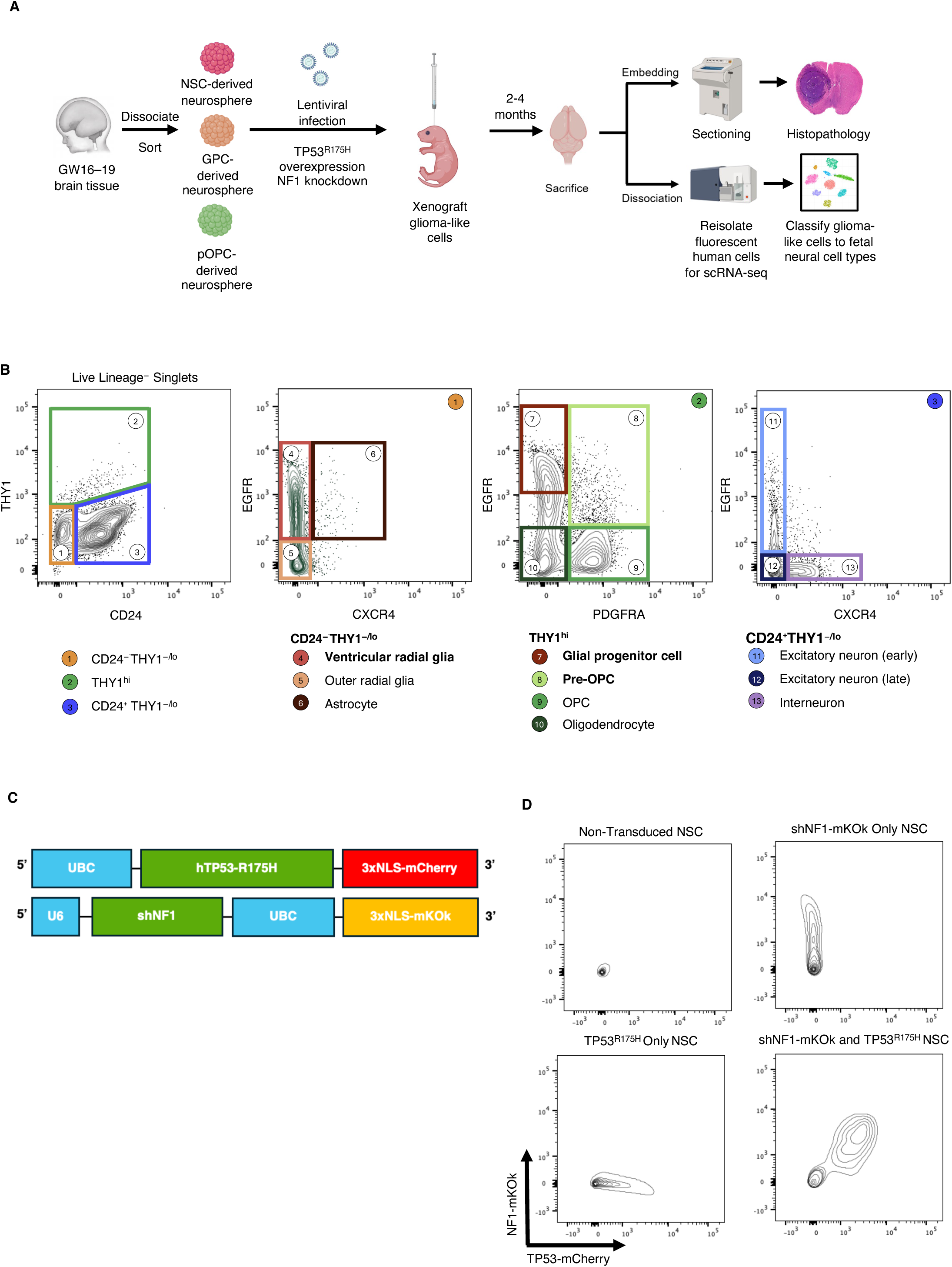
Workflow of transformed NSPCs. **a.** Experimental workflow used to collect histological samples and dissociated cells from mice injected with transformed NSPCs. **b.** FACS sort scheme to collect isolated NSPCs from fetal human brain. Adapted from Liu et al. 2023. Lineage exclusion included antibodies against CD31, CD34, CD45, CD105, and CD235a. **c.** Lentiviral constructs used for TP53^R175H^-mCherry and shNF1-mKOk double knockout **d.** FACS sort scheme to collect mCherry and mKOk double positive cells

### TP53^R175H^/shNF1-NSPCs engraft, migrate, and form disorganized lesions

To evaluate the engraftment and migration of our TP53^R175H^/shNF1-NSPC lines, we transplanted them into the brain parenchyma of neonatal NOD scid gamma (NSG) mice. Cells were transplanted unilaterally in the left hemisphere to assess migration. Mice were sacrificed at 2 and 4 month time points, after which brains were taken for either histology or single cell RNA-seq of engrafted human cells. Confocal immunofluorescence microscopy showed that following transplant, TP53^R175H^/shNF1-NSPC populations robustly engrafted **(Figure 2A)**. Furthermore, we noticed disorganized lesions of human cells **(Figure 2B)**. Strikingly, TP53^R175H^/shNF1-NSPCs aggressively migrated to the contralateral hemisphere through the corpus callosum, reminiscent of invasive “butterfly” gliomas **(Figure 2C)**^22^. In contrast, wild-type NSCs that did not receive the TP53^R175H^ or shNF1 constructs remained localized at the transplant site, with no cells found crossing the midline **(Figure 2D)**.

**Figure 2:**
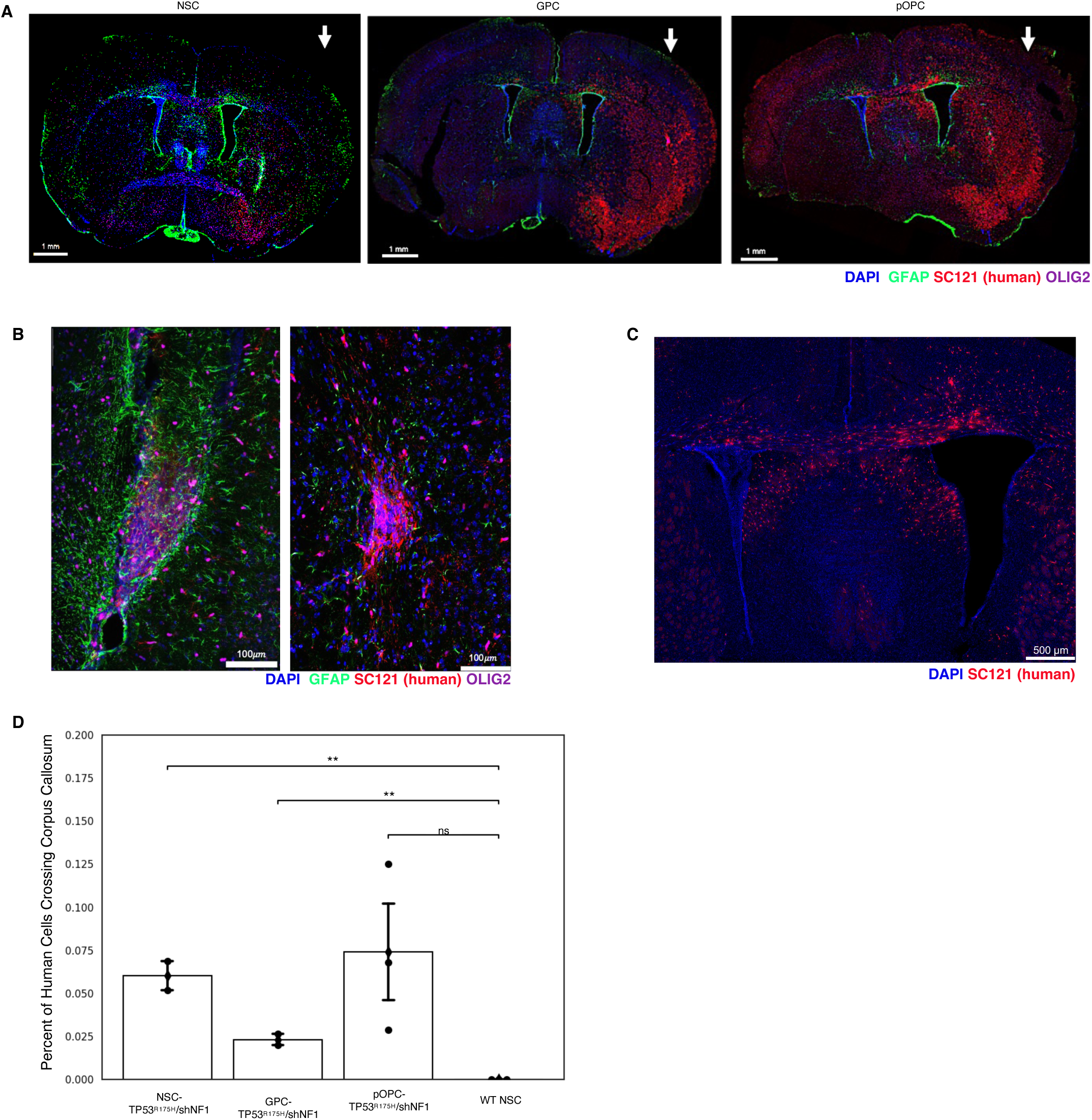
Flow cytometry results of transformed NSPCs collected after 2 months and 4 months. **a.** Image of whole brain of mice injected with transformed NSPCs. Arrow indicates the hemisphere of transduced NSPC injection. **b.** Zoomed-in image of brain injected with transformed NSCs to show lesion formation **c.** Zoomed-in image of brain injected with transformed pOPCs to show corpus callosum migration **d.** Quantification of migrating NSPC-derived cells crossing the corpus callosum

### TP53^R175H^/shNF1-NSPCs expand over time following transplantation

In addition to histology, we dissociated mouse brain tissue engrafted with TP53^R175H^/shNF1-NSPCs at 2- and 4-month time points and analyzed with flow cytometry. Engrafted human cells were identified based on their mCherry and mKOk reporters. The proportion of engrafted human cells increased 6-fold between the 2- and 4-month time point, suggesting prolonged expansion post-transplant **(Figure 3A, B)**.

**Figure 3:**
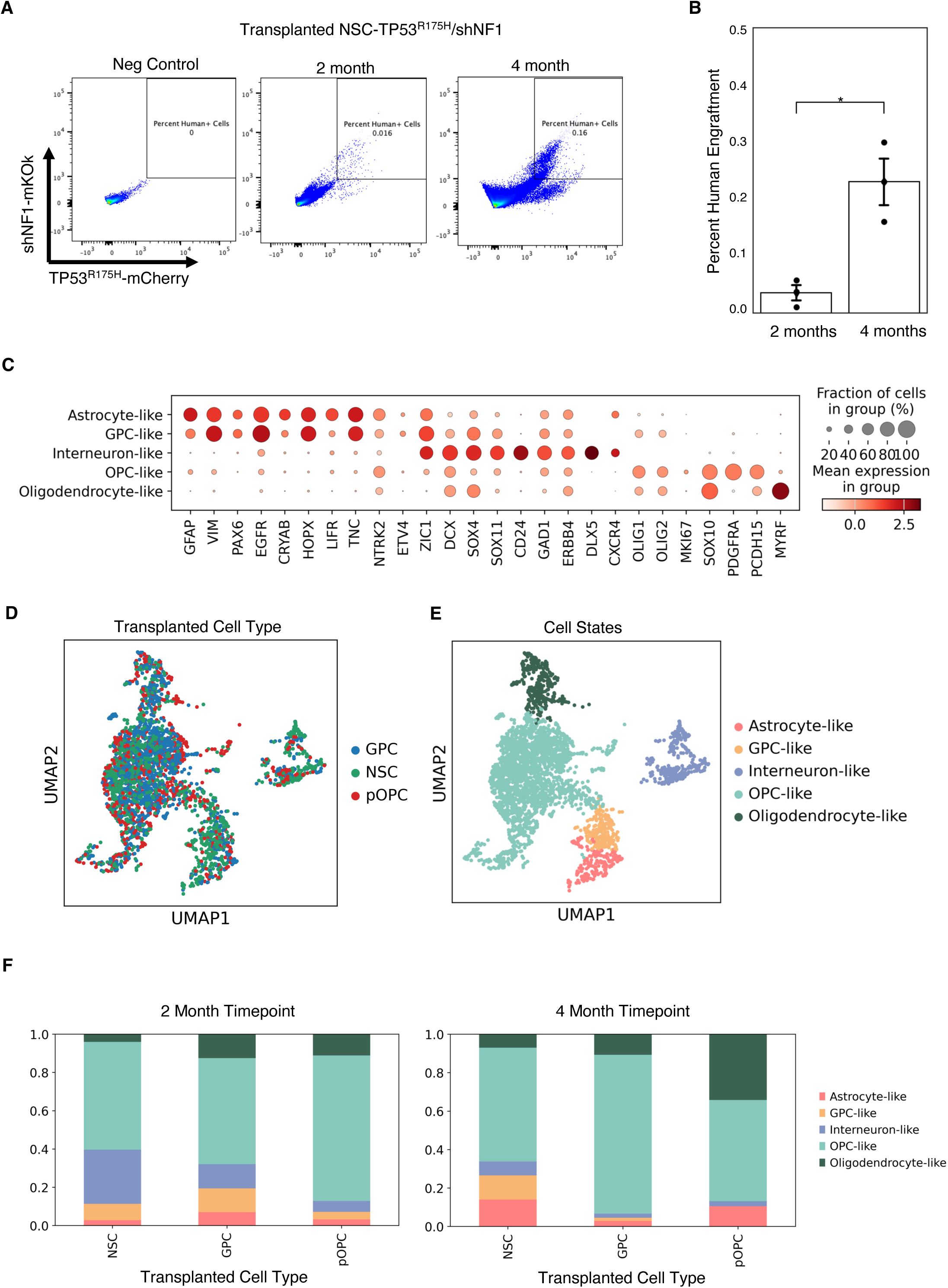
Single cell RNA sequencing analysis of transformed NSPCs. **a.** FACS plots of mice after 2 and 4 months of transplantation, with an uninjected mouse for comparison **b.** Quantification of human cells collected from NSPC conditions between 2 month and 4 month timepoints **c.** Dot plot of distinguishing markers used to identify xenografted cell identity **d.** UMAP plot of xenografted NSPCs annotated by NSPC identity **e.** UMAP plot of xenografted NSPCs annotated by expression-based cell identity **f.** Stacked bar graph showing percentages of cell type composition in each injected NSPC condition at the 2 month takedown (left) and 4 month takedown (right).

### TP53^R175H^/shNF1-NSPCs recapitulate glioma intratumoral heterogeneity

To assess the fate of our TP53^R175H^/shNF1-NSPC lines following transplant, we collected engrafted cells at the 2 and 4 month timepoints for single cell RNA-seq (scRNA-seq) using a modified Smart-seq3 protocol^23^. The engrafted cells recapitulated all major neural lineages, including clusters resembling OPCs, oligodendrocytes, astrocytes, GPCs, and neurons **(Figure 3C)**. There were no major differences in the transcriptomic profiles of the engrafted cells at 2 months vs 4 months **(Supplementary Figure 1)**. Interestingly, at 2 months post-transplant, all three isogenic lines (NSC-, GPC-, and pOPC-derived) gave rise to all neural lineages **(Figure 3D–F)**, including GPC-, OPC-, oligodendrocyte-, astrocyte-, and interneuron-like cells. However, at 4 months the GPC and pOPC transfected cells gave rise to predominantly glial lineages.

This is in contrast with our previous studies using acutely-isolated NSPC populations, wherein GPCs were lineage restricted to astrocytes and oligodendrocytes, and OPCs gave rise exclusively to oligodendrocytes^18^. However, the relative makeup of engrafted cells still differed based on the parental cell line **(Figure 3F)**. NSC-derived cells gave rise to the highest proportion of neuronal-like cells (28%). pOPC-derived cells, despite minor contributions to other lineages, still gave rise nearly exclusively to oligodendrocyte lineages (87%), and resulted in the highest percentage of differentiated oligodendrocyte-like cells at 4 months post-transplant (34%). These differences suggest that cell-of-origin does impact the relative makeup of resulting tumors, though the lineage restriction may be less strict than in normal development.

## DISCUSSION

Here we have shown that purified fetal human NSPCs transduced with driver mutations can serve as a robust cellular model for glioma heterogeneity. Transformation of purified NSPC populations with TP53^R175H^ and shNF1 endowed cells with tumorigenic and invasive properties. The single cell analysis demonstrates that the cell-of-origin (NSC vs. GPC vs. pOPC) influences the relative makeup of resulting glioma-like cells. Together, these results suggest that glioma initiation mirrors normal neural stem cell development, and the methods demonstrate that using purified fetal neural stem and progenitor cells are a useful model for gliomas of varying extents of intratumoral heterogeneity.

There are of course various mouse models of glioma that similarly induce driver mutations in select subsets of neural cells, usually via knockout of TP53, NF1, and PTEN^24–26^. These studies have shown that such knockouts in either NSCs or OPCs were sufficient to induce tumors, though the cell-of-origin did seem to impact the subtype of the resulting glioma, with OPC-derived gliomas largely expressing oligodendrocyte-lineage GFAP^low^ tumor cells and NSC-derived gliomas expressing both GFAP^high^ and GFAP^low^ tumor cells^27^.

Our model system, using reprogrammed primary human fetal NSPCs, holds a few distinct advantages over mouse models. First, it is highly modular, allowing for the introduction of specific genetic manipulations into specific isogenic cell types at defined developmental stages (NSC, GPC, pOPC) with a high turnaround rate. Second, it more faithfully models human neurodevelopment programs, which has important differences compared to that of mice. For example, the presence of oRG and GPC cell types is much less well-defined in murine corticogenesis^28^. Third, because these lines can be derived from various fetal tissue donors, this model system allows investigation of how different genetic backgrounds impact glioma initiation as well.

It has long been observed that glioma cells mirror normal neural cell types. Single cell transcriptomics has now enabled resolution of this intratumoral heterogeneity. Notably, gliomas of different histological subtypes and grades demonstrate different admixtures in terms of cellular composition^10^. IDH-mutant oligodendrogliomas are enriched for differentiated oligodendrocyte-like cells, whereas H3K27M gliomas are enriched in more proliferative OPC-like cells. IDH-wildtype glioblastomas harbor an even more diverse repertoire of cellular subtypes, including a neuronal subtype endowed with the remarkable migratory capabilities characteristic of newborn cortical neurons. These observations suggest the presence of a stem cell hierarchy within gliomas that recapitulate fetal neurodevelopment. Our study demonstrates that the cell-of-origin impacts the downstream cellular makeup of resulting gliomas. Thus, different glioma types may be better modeled with different NSPC lines; for instance, TP53^R175H^/shNF1-NSCs may best model GBM, given their ability to robustly generate a neuronal-like cellular subtype. We also observe that TP53^R175H^/shNF1-pOPCs have a propensity to produce more differentiated oligodendrocyte-like cells, whereas TP53^R175H^/shNF1-GPCs primarily produce more OPC-like cells; it is tempting to draw a parallel here with the lower grade IDH-mutant oligodendrogliomas versus the higher grade H3K27M gliomas.

Future work using our platform may more deeply explore the impact of other genetic perturbations on resulting glioma heterogeneity, for example, ATRX, EGFR, PDGFRA, and CDK4. More than just a tool for interrogating intratumoral heterogeneity, these lines may also be used to test the efficacy of therapeutic interventions based on cell-of-origin and genetic drivers. Of course, all models still have limitations. In our case, the artificial introduction of TP53^R175H^ and NF1 genetic perturbations may not accurately reflect the natural history of glioma evolution in patients. Indeed, TP53 is often the final mutation in other cancer types, most famously the adenoma-carcinoma sequence in colon cancer^29^. It is also possible, if not likely, that the gene regulatory network present in fetal NSPCs does not accurately reflect those in adult NSPC cells, though we still believe these cells are the best option given that prospective isolation of viable NSCs from adult human brains has not yet been achieved. We posit that our reprogrammed NSPCs present a useful and modular platform for investigating cell-of-origin impacts on glioma intratumoral heterogeneity.

## ACKNOWLEDGMENTS

We thank Teja Naik, Linda Quinn for laboratory management; Catherine Carswell-Crumpton and Cheng Pan for flow cytometry support; Katia Alvarez, Michael R. Eckart, and the Stanford Protein and Nucleic Acid (PAN) Facility for technical support with cDNA quality control; Norma Neff, Honey Mekonen, Angela Detweiler, and the Chan Zuckerberg Biohub for sequencing support. D.D.L is supported by Stanford University Medical Scientist Training Program grant T32-GM007365 and T32-GM145402, and the Seth A. Ritch Bio-X Stanford Interdisciplinary Graduate Fellowship (SIGF). This work was supported by an NIH/NCI Outstanding Investigator Award R35-CA220434 and the Virginia and D.K. Ludwig Fund for Cancer Research to I.L.W.

## AUTHOR CONTRIBUTIONS

D.G., D.D.L., A.E.E., and N.L.W. performed the experiments. D.G., B.F.O.-G., D.D.L., and M.B. processed Smart-seq3 libraries. D.D.L. and D.G. analyzed the data. D.D.L. and D.G. wrote the manuscript. I.L.W. and D.D.L. supervised the study. All authors approved the final manuscript.

## METHODS

### Mice

Fetal human NSPCs were xenografted into NOD.Cg-Prkdc^scid^ Il2rg^tm1Wjl^/SzJ (NSG) immunodeficient mice (JAX: 005557). These mice were housed in Stanford University’s animal core facility following guidelines set by the National Institutes of Health (NIH) and Stanford’s Administrative Panel on Laboratory Animal Care (APLAC).

### Fetal brain samples

Deidentified human fetal brain sample were obtained through Advanced Bioscience Resources (ABR, Alameda, CA). Collected sample ages ranged between 16 and 19 gestational weeks. All experiments using this tissue were performed under guidelines set by Stanford University’s Stem Cell Research Oversight (SCRO) (SCRO protocol #735, APLAC protocol #26209). No Institutional Review Board (IRB) approval was required for deidentified samples.

### Fetal brain dissociation

Samples were dissociated into a single cell suspension as previously described^18,20^. In brief, samples were chopped with a razor blade and resuspended in 10 μg/mL Liberase (Roche, 5401119001) and 200 μg/mL DNase I (Worthington, LS002007). Then the samples were incubated on a shaker at 37°C for 30 minutes and triturated through a P1000 pipette tip attached to a serological pipette. Samples were then resuspended in Accutase (Innovative Cell Technologies, AT104) with 200 μg/mL DNAse I, then incubated again on a shaker at 25°C for 15 minutes. Red blood cells were removed by performing a density gradient spin with Histopaque (Sigma, 10771) at a 2:1 ratio in a centrifuge set at 400×g for 30 minutes at 25°C with brakes off. The buffy coat was collected after the spin to be washed, counted, and stained for flow cytometry and cell sorting.

### Isolation of human NSPCs

Dissociated human fetal brain samples were stained with antibodies against CD24 (Clone 32D12, Miltenyi, 130095954), CD90 (Clone 5E10, Biolegend, 328108), CXCR4 (Clone 12G5, Biolegend, 306528), EGFR (Clone AY13, Biolegend, 352910), PDGFRA (Clone 16A1, Biolegend, 323504), CD31 (Clone WM59, BD Biosciences, 563652), CD34 (Clone 581, BD Biosciences, 562383), CD45 (Clone HI30, Biolegend, 304010), CD105 (Clone 266, BD Biosciences, 562380), and CD235a (Clone HIR2, Biolegend, 306606). All antibodies were diluted at a ratio of 1:50. Propidium iodide (PI) was added immediately before flow analysis to a final concentration of 1 μg/mL. Flow cytometry and fluorescence-activated cell sorting (FACS) utilized a FACS Aria II (BD Biosciences) for analysis. Gates were drawn to exclude debris, doublets, dead cells and hematopoietic lineages. Cells were collected using the “4-way purity” setting for bulk sort.

### Cell Culture

All cells were cultured in Fetal Growth Media (FGM)^20^. Cells were initially suspended in 200,000 cells/mL. After 5 days, FGM was added such that the final cellular density was 100,000 cells/mL, calculated from the Day 0 cell count. After 10 days, cells were dissociated with Accutase (Innovative Cell Technologies, AT104) in preparation for viral transduction or xenotransplantation.

### Lentivirus Production and Infection

Lentivirus for NF1 shRNA (sense, 5′-CUGUGUAAAGCAAGUACUU-3′) and TP53^R175H^ (anti-sense, 5’-CGC-3’ to 5’-CAC-3’ at nucleotide positions 522-524) were produced by first combining 11 μg of lentivirus construct with 5.5 μg of pMD2.G (Addgene, #12259) and 8.25 μg of psPAX2 (Addgene, #12260) in 1.5 mL of OptiMEM and mixed. Then, 74.5 μL of Trans-IT (Mirius Bio, MIR 2700) was added and slowly mixed with a P1000 pipette. The mixture was incubated for 20 min at room temperature before being added to 293T cells fed with OptiMEM media and incubated at 37°C. After 24 hours (Day 1), the 293T media was collected and stored at 4°C and the cells refreshed with 7 mL of OptiMEM. After another 24 hours (Day 2), the media was again collected and pooled with the Day 1 media and filtered through a 0.45 μm filter. The virus was precipitated by adding 50% polyethylene glycol (PEG) at ¼ the volume of the total virus media, vortexed, then stored at 4°C for 24 hours. After, the virus was centrifuged at 2,500G for 20 min with bucket caps to pellet, and the supernatant was aspirated. The pellet was resuspended in 100 uL of HBSS. Virus was titrated to achieve a 30-50% efficiency. Single-cell suspension of NSPCs were incubated with virus for 90 min at 37°C on a shaker. After, cells were recultured in FGM as previously described above.

### NSPC Transplantation

NSPCs were resuspended in a mixture of HBSS and Fast Green dye (Sigma, F7252) to a final concentration of 10^5^ cells/1 μL. Neonatal mice (postnatal day 1-3) were anesthetized on ice for 2 minutes and positioned on a stereotactic device (Harvard Apparatus). Two burr holes were made in the left parenchyma by a 30-gauge needle to identify the injection sites. Cells were injected into mice by a 33-gauge Hamilton syringe controlled by a Micro4 microsyringe pump (World Precision Instruments). Cells were injected at a rate of 1 μL/min and 0.5 μL was injected per site, for a final 10^5^ cells transplanted per mouse.

### Mouse Brain Dissociation

Mice were sacrificed 2 months and 4 months post-transplant. Brain tissue was collected and chopped with a razor blade to ∼1 mm^3^ pieces. The chopped tissue was added to Papain (Worthington, LK003178) and 50 μg/mL DNase I (Worthington, LS002007) and placed on a shaker at 37°C for 20 minutes at 750 RPM. Samples were then pulled through a 1000 μL pipette tip attached to a serological pipette until the cell suspension could pass through smoothly, and placed back on the shaker for an additional 10 minutes. The cell suspension was filtered through a 70 μm cell strainer. Myelin debris was then depleted by density gradient. First, 4 mL of Debris Removal Solution (Miltenyi, 130-109-398) was added then overlayed with 4 mL of cold HBSS, forming two phases. This suspension was spun down at 3000G for 10 min. Then, the resulting top two phases were transferred to the Percoll density gradient (Cytiva, 17-0891-02) while the bottom phase containing the cell pellet was washed with HBSS and placed on ice. Cells were added to a tube containing 20% Percoll and centrifuged at 300×g at 25°C for 15 min with brakes turned off. The myelin layer and supernatant were removed, and the cell pellet was recombined with the cell pellet in the Debris Removal Solution Step. This pellet was then resuspended in 1 mL ACK lysis Buffer (Gibco, A1049201) for 2 min. 9 mL of HBSS was added to stop the lysis action. The dissociated samples were stained with antibodies against mouse CD45 (clone 30-F11, Biolegend, 103110) and Ter119 (clone TER-119, Biolegend, 116209) for FACS. SYTOX Red (ThermoFisher Scientific, S34859) was added at a volume of 1 μg/mL before analysis for measuring viability. Live singlet events that were positive for mKOK and mCherry and negative for CD45 and Ter119 were isolated using FACS and taken for Smart-Seq3 sequencing.

### Histology

Mice were anesthetized using isoflurane and perfused with cold PBS with 5 mM EDTA (ThermoFisher Scientific, 15-575-020) through the left ventricle. Mouse brains were dissected out and fixed in 4% paraformaldehyde (PFA) (Electron Microscopy Sciences, 15710) for 24 hours, then transferred to 30% sucrose solution for 24 hours. Coronal sections were sliced 40 μm thick using a Leica SM2010 R Sliding Microtome. Tissue slices were kept in Tissue Collection Solution (25% glycerin, 30% ethylene glycol in phosphate buffer) at -20°C before staining.

### Immunofluorescence

Coronal brain sections were blocked and permeabilized in 1X TBS Buffer (ThermoFisher Scientific, 28358) with 0.3% Triton and 3% normal horse serum (ThermoFisher Scientific, MT35030CV) for 1 hour. Then, sections were stained overnight in 4°C with primary antibodies: human cytoplasmic antigen (1:1000, clone STEM121, Takara, Y40410), OLIG2 (1:1000, AbCam, 109186), GFAP (1:1000, NovusBio, 05198). Then, sections were washed thrice with 1X TBS Buffer, then incubated in secondary antibodies for 2 hours (ThermoFisher Scientific). Primary and secondary antibodies were diluted in 1X TBS with 0.3% Triton and 1% normal horse serum. Then, samples were placed in DAPI (1 μg/mL) for 10 min and washed thrice with 1X TBS Buffer. Samples were then mounted on glass slides using Prolong Gold (ThermoFisher Scientific, P36930) and imaged on a Leica Mica confocal microscope. For quantification of cells crossing the corpus callosum, sections of the brain from Bregma 1.09 mm to -2.53 mm were used. Cells positive for SC121 were manually counted, and the percentage of human cells crossing the midline was calculated for each section. Significance was calculated using the student t-test.

### Single Cell Sequencing

Collected cells for single cell were processed using a modified protocol of Smart-seq 3^20,23^. Live singlet events that were double-positive for TP53^R175H^-mCherry and shNF1-mKOK were single-cell sorted into 96 well plates. Each well contained 2 μL lysis buffer (0.1% Triton (Thermo Fisher Scientific, 85111), 2.5 mM dNTP (Thermo Fisher Scientific, 10297018), 1 U/μL RNase inhibitor (Clontech, 2313B), 2.5 μM oligo dT_30_VN (Integrated DNA Technologies, 5′-AAGCAGTGGTATCAACGCAGAGTACT_30_VN-3’), and 1:600,000 ERCC RNA spike-in mix at 1:600,000 (Thermo Fisher Scientific, 4456739) in UltraPure water (Thermo Fisher Scientific, 10977015)). Immediately after a completed collection, plates were snap-frozen on dry ice and stored at −80°C.

When preparing the cells for reverse transcription (RT), the plates were incubated at 72°C for 3 minutes and immediately snap-chilled on ice for primer annealing. Then, 3 μL RT mix was added to each well. This mix contained 25 mM Tris-HCl pH 8.5 (Teknova, T5085), 30 mM NaCl (Thermo Fisher Scientific, AM9760G), 2.5 mM MgCl_2_ (Thermo Fisher Scientific, AM9530G), 8 mM dithiothreitol (DTT) (Promega, P1171), 0.5 U/μL RNase inhibitor (Clontech, 2313B), 1 mM GTP (Thermo Fisher Scientific, R0461), 2 μM TSO (Integrated DNA Technologies, 5′-AAGCAGTGGTATCAACGCAGAGTGAATrGrGrG-3′), 5% polyethylene glycol (Sigma, P1458), and 2 U/μL Maxima H-minus reverse transcriptase (Thermo Fisher Scientific, EP0753) in UltraPure water. After RT mix was added, plates were incubated at 42°C for 90 min, then at 70°C for 15 min. Then 7 μL PCR mix was added to each well to amplify cDNA content. The PCR mix contained 1.67X KAPA HiFi HotStart ReadyMix (Kapa Biosystems, KK2602) and 0.17 μM ISPCR primer (Integrated DNA Technologies, 5′-AAGCAGTGGTATCAACGCAGAGT-3′) in UltraPure water. Then, the following program was applied to the samples: (1) 98°C for 3 min, (2) 98°C for 20 sec, (3) 67°C for 15 sec, (4) 72°C for 6 minutes, (5) repeat from Step 2 24 times, and finally (6) 72°C for 5 minutes. To purify cDNA, the amplified cDNA was cleaned with 0.65X volume of calibrated AMPure XP beads (Beckman Coulter, A63882), washed twice with 80% ethanol, then eluted in 12.5 μL UltraPure water.

1· μL of purified cDNA was taken on a capillary electrophoresis-based Fragment Analyzer (Advanced Analytical) for quantity control and cDNA concentration assessment. Cells with a concentration less than 0.5 ng/μL were excluded. The remaining cells were transferred into 384 well plates using a Mosquito X1 liquid handler (SPT Labtech). During this process, each cell’s concentration was normalized by diluting with UltraPure water for a final concentration range of 0.5-4 ng/μL.

0.4 μL of normalized cDNA was added to 1.2 μL homebrew Tn5 mix (1 ng/μL Tn5 enzyme, 16 mM Tris-HCl pH 7.6, 16 mM MgCl_2_ (Thermo Fisher Scientific, AM9530G), 8% dimethylformamide (DMF) (Thermo Fisher Scientific, AC327171000) in UltraPure water) in 384 well plates and incubated at 55°C for 10 min. Then, 0.4 μL neutralization buffer (0.1% SDS) was added to stop the reaction. To index and amplify the Tn5 products, 0.4 μL of 5 μM i5 indexing primer, 0.4 μL of 5 μM i7 indexing primer (Integrated DNA Technologies, custom made 7680-plex unique dual index-primer set), and 1.2 μL KAPA HiFi HotStart ReadyMix were added to each well. Then, the following incubation program was run: (1) 72°C for 3 min, (2) 95°C for 30 sec, (3) 98°C for 10 sec, (4) 67°C for 30 sec, (5) 72°C for 60 sec, (6) repeat from step 3 10 times. Each 384 well plate was then pooled into a 1 μL reaction product and cDNA was purified using 0.8X volume of AMPure XP beads (Beckman Coulter, A63882). Each pool was analyzed for concentration and size distribution using Bioanalyzer (Agilent).

Reads were demultiplexed using bcl2fastq v2.20. 3’ adapter sequences were removed from reads with skewer v0.2.2, then aligned to the hg38 genome (GRCh38.p13) with STAR v2.6.1d using 2-pass mapping. For the first pass, reads for every cell were aligned using a STAR genome index that was generated with the Gencode transcript annotation for the human genome (v34). Splice junctions that resulted from this first pass were extracted and aggregated to generate a custom STAR genome index, which includes newly discovered splice junctions in addition to the existing Gencode annotations. This custom genome index was used for second-pass mapping. STAR mapping parameters were adapted from the ENCODE long-mRNA-pipeline recommendations. In addition, “--quantMode TranscriptomeSAM” was added during the second pass mapping to generate a bam file that catalogs expression levels of either the total sum of expression levels of all known transcript variants (“genes”) or individual transcripts. Gene and individual transcript counts were tabulated with RSEM v1.3.3 with the “--single-cell-prior” setting.

---

**Supplemental Figure 1:**
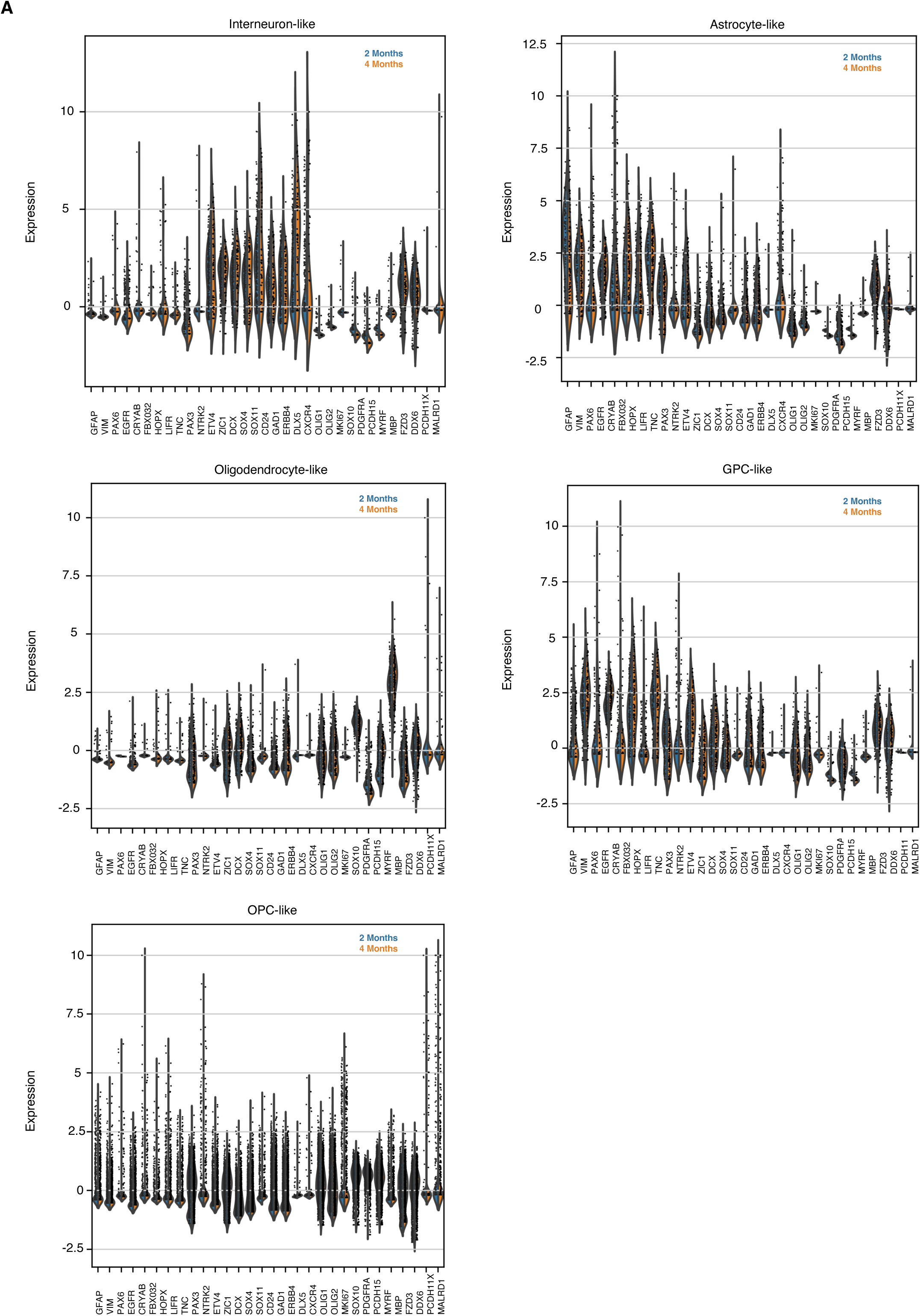
Differential Gene Analysis of Cell Types between 2 Month and 4 Month Time Points. **a.** Violin plots of cell types and gene expression between 2 month and 4 month timepoints

